# Mitochondrial RNA granules are fluid condensates, positioned by membrane dynamics

**DOI:** 10.1101/747055

**Authors:** Timo Rey, Sofia Zaganelli, Emilie Cuillery, Jean-Claude Martinou, Suliana Manley

## Abstract

Mitochondria contain the genetic information and expression machinery to produce proteins essential for cellular respiration. Within the mitochondrial matrix, newly synthesized RNA, RNA processing proteins, and mitoribosome assembly factors are known to form punctate subcompartments referred to as mitochondrial RNA granules (MRGs) ^1–3^. Despite their proposed role in regulating gene expression, little is known about the structural and dynamic properties of MRGs. We investigated the organization of MRGs using fluorescence super-resolution localization microscopy and correlative electron microscopy techniques, obtaining ultrastructural details of their internal architecture. We find that MRGs are organized into nanoscale RNA cores surrounded by a protein shell. Using live-cell super-resolution structured illumination microscopy and photobleaching perturbations, we reveal that MRGs undergo fusion and rapidly exchange components, consistent with liquid-liquid phase separation (LLPS). Furthermore, MRGs associate with the inner mitochondrial membrane and their fusion coincides with membrane remodeling. Inhibition of mitochondrial fission leads to an aberrant distribution of MRGs into concentrated pockets, where they remain as distinct individual units despite their close apposition. Together, our results reveal a role for LLPS in concentrating RNA and its processing proteins into MRGs, which are positioned along mitochondria by membrane dynamics.

---

RNA in eukaryotic and bacterial cells is sequestered into granules that exhibit a wide range of forms and functions, under both physiological and stress conditions. For example, nuclear speckles and paraspeckles, are involved in RNA splicing and transcriptional regulation ^4–6^. The nucleolus creates a compartment for ribosomal assembly ^7, 8^, and in the cytoplasm, stress granules protect mRNAs during cellular stress ^9^. RNA-protein granules often form by liquid-liquid phase separation ^10^. In developing embryos, the sensitivity of P granule formation by LLPS to the local concentration of proteins leads to symmetry breaking and differential inheritance of material between germ and somatic cells ^11^. Multivalent weak interactions between disordered RNA-binding protein (RBP) domains, and RNA itself, were identified as crucial factors for the formation of biomolecular condensates in many *in vitro* and *in silico* studies ^12–15^. To understand their biological function, *in vivo* studies are essential; phase behavior is sensitive to elements of the molecular environment such as salt concentration, pH, or crowding, and physiological conditions are challenging to reproduce in test tubes ^16^.

MRGs are ribonucleoprotein complexes composed of newly synthesized long polycistronic mtRNAs, originating from the 16kb mitochondrial DNA (mtDNA), together with several mitochondrial RBPs ^1, 17, 18^. It was previously demonstrated that mtRNA is essential for MRG formation ^3^. However, both the structural organization of and the dynamic interplay between MRG components are still unknown. Mitochondria undergo dramatic shape changes through fission, fusion, and branching ^19^, of which fission directly impacts the distribution of mtDNA ^20, 21^. How MRGs respond to this dynamicity and complex architecture is also unknown, due in part to their size below the diffraction limit. Here, we investigated the molecular organization, distribution, and positioning mechanism of MRGs within the mitochondrial network, using super-resolution and correlative electron microscopy. We show MRGs are ∼130 nm, sub-compartmentalized liquid condensates, associated with the inner mitochondrial membrane (IMM), which mislocalize upon perturbation of mitochondrial fission.

First, to assess MRG dimensions and overall organization, we immunolabeled MRG-associated RBPs in fixed mammalian cells. We chose two *bona fide* MRG markers, GRSF1 and FASTKD2, for which mutations associate with severe mitochondrial diseases ^22, 23^. In conventional fluorescence images of FASTKD2, MRGs appear as bright puncta against a dim mitochondrial matrix filled with free FASTKD2 (Fig. 1a). We used reference fluorescence images to identify MRGs, and the correlate high throughput stochastic optical reconstruction microscopy (htSTORM) images to compute their size with a precision down to 10 nm ^24^ (Supplementary Fig. 1). We found similar median diameters for the two markers: 137 nm (± 32 (SD), *n* = 326) for FASTKD2, and 122 nm (± 30 (standard deviation, SD), *n* = 361) for GRSF1 foci in COS-7 cells (Fig. 1a, b **and** Supplementary Fig. 2, 3). To assess the arrangement of RNA components of MRGs we incubated the cells with 5 mM bromouridine (BrU) for one hour and immunolabelled the newly synthesized mtRNA with anti-BrU as originally described ^1, 3^. mtRNA distributed within MRGs occupy a region with a median diameter of 94 nm (± 49 (SD), *n* = 254) (Fig. 1b **and Supplementary Fig. 4**), on average significantly smaller than volumes occupied by the protein markers, albeit with a larger variance. Nucleoids containing mtDNA also constitute punctate structures ∼100 nm in radius ^25–27^, serving as an internal reference. In good agreement with reported values, we observed a median diameter of 91 nm (± 36 (SD), *n* = 431) for antibody-stained mtDNA (Fig 1b **and Supplementary Fig. 5**). Next, we performed two-color htSTORM to understand how mtRNA and FASTKD2 organize within MRGs. We found that many mtRNA foci appear to be encased by protein (Fig. 1c). Altogether, this shows that MRGs have an internal structure of a multi-component protein shell surrounding one or more RNA cores, as proposed for stress granules^28^.

**Fig. 1.:**
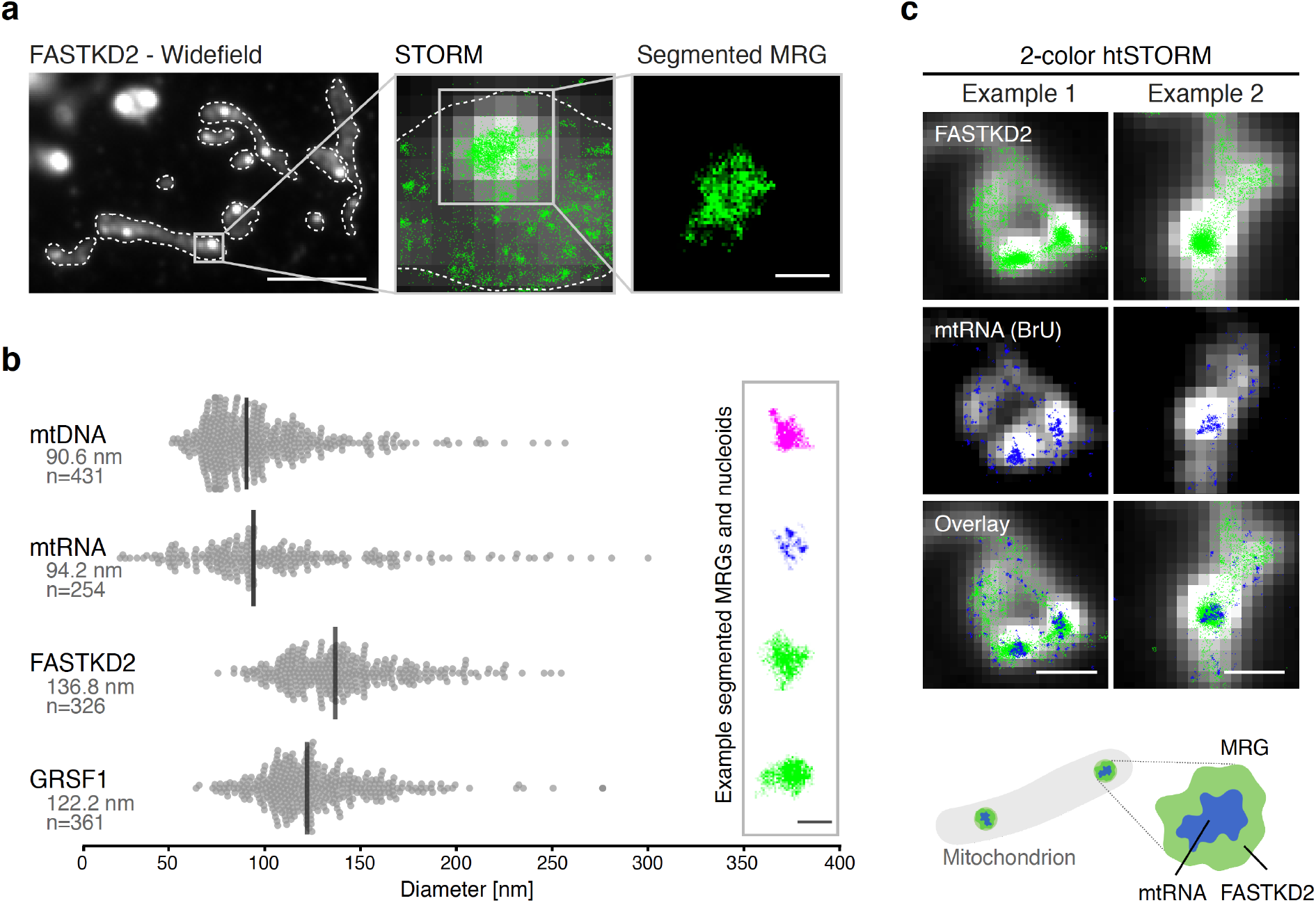
Nanoscale organization of MRGs. **a**, MRG size assessment by super-resolved htSTORM. Widefield immunofluorescence of FASTKD2 shows punctate MRGs within mitochondria (white dashed lines). Scale bar: 10 µm. Zooming in, a STORM image (green) shows a single MRG, overlaid on a widefield image (greyscale). The MRG is segmented by its high density of STORM localizations using DBSCAN. Scale bar: 200 nm. **b**, MRG and nucleoid (mtDNA) diameters measured from DBSCAN-clusters. MRGs were imaged using different markers (mtRNA, FASTKD2 and GRSF1). Median diameter and number of analyzed clusters per condition are noted. Data points are shown in grey, median is indicated as black line. A representative image for each condition is shown on the right. Scale bar: 200 nm. **c**, Examples of 2-color htSTORM of anti-FASTKD2 (MRGs, green) and anti-BrdU (mtRNA, blue). htSTORM images are overlaid on widefield images (grey). Scale bar: 500 nm.

Liquid-phase properties of stress-and other RNA granules play a role in mRNA sequestration, enzyme buffering, and tuning of reaction kinetics ^10^. These roles may apply to MRGs and their function in gene expression, if LLPS underlies MRG formation. To test this hypothesis, we generated stable FASTKD2-eGFP expressing cell lines and assessed common hallmarks for cellular LLPS: content exchange and droplet fusion ^11^. We examined the molecular exchange of MRG components by fluorescence recovery after photobleaching (FRAP). To monitor MRG fluorescence inside highly mobile mitochondria, we developed a software tool for FRAP analysis with motion tracking. We found FASTKD2-eGFP molecules within MRGs to recover rapidly in both U2OS and COS-7 cells, with a half-recovery time of 6.5 seconds (Fig. 2a, c **and Supplementary Fig. 6a-d**). To test whether dynamic exchange is generalizable beyond FASTKD2, we created two additional stable cell lines expressing MRG markers, ERAL1 and DDX28, fused to eGFP ^29, 30^. Both recovered rapidly, at a timescale (4.4 seconds, ERAL1 and 6.3 seconds, DDX28) similar to FASTKD2 (Fig. 2c). For comparison, we overexpressed the mitochondrial helicase TWINKLE fused to eGFP as a nucleoid marker with high DNA-binding affinity ^31^. As expected, TWINKLE foci only slightly recover over the course of our FRAP assay (50 sec) (Fig. 2b, d). Thus, MRG components exchange rapidly, on a fast timescale compared to stress granules ^32^.

**Fig. 2.:**
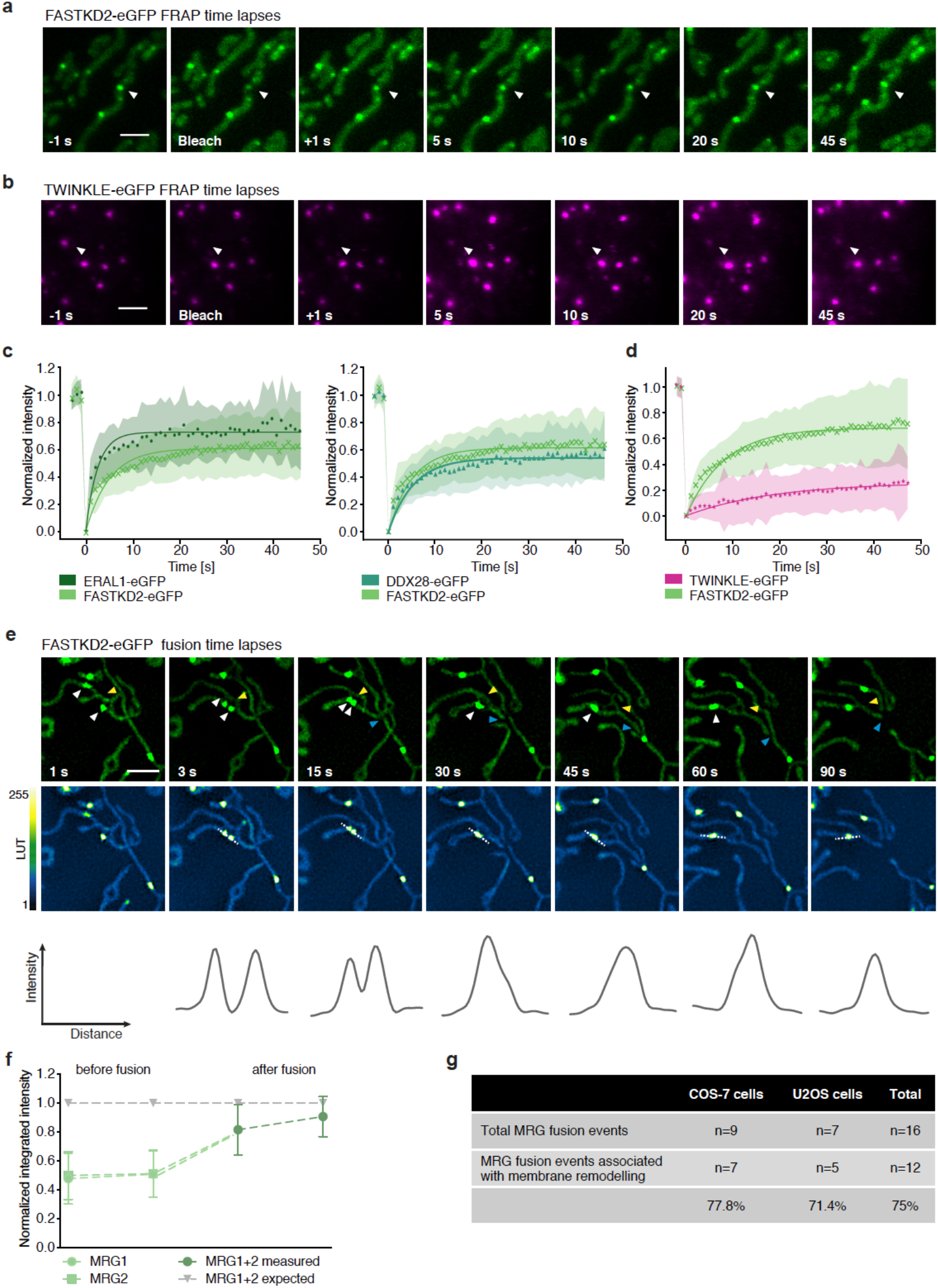
Content exchange and fusion of MRGs in live COS-7 cells. **a, b**, Representative FRAP time lapse images in cells stably expressing FASTKD2-(MRGs, green) or TWINKLE-eGFP (nucleoids, magenta). Arrowheads indicate the target structures, partially photobleached to allow tracking. Scale bars: 2 µm. **c, d**, FRAP-recovery plots and analysis for FASTKD2-, ERAL1-, DDX28-(**c**) and TWINKLE-eGFP (**d**). Symbols in the graph represent mean data points. Single exponential fits (lines) and standard deviations for each time point (shaded areas) are shown **e**, Representative SIM time lapse images of an MRG fusion event in cells stably expressing FASTKD2-eGFP. Cells were imaged at 1/3 Hz. White arrowheads indicate the MRGs fusion event, yellow and blue arrowheads highlight mitochondrial dynamics. The graphs show the fluorescence intensity (pixel values) along the dashed lines at each time point. Scale bar: 2 µm. **f**, Temporal evolution of the integrated intensity of MRGs in COS7 before and after fusion. Pre-fusion intensities were normalized by their sum for each granule-pair (n=9). Mean and SD for each MRG type are shown for two timepoints before as well as after fusion. **g**, Summary of MRG fusion events observed, and associated with mitochondrial membrane remodeling.

To determine whether MRGs were better described as liquid droplets or solid granules, we followed FASTKD2 foci by live-cell super-resolved structured illumination microscopy (SIM). The resolution of SIM is sufficient to discern fusion of liquid drops from aggregation of solid granules. We observed MRG fusion in multiple instances in both U2OS and COS-7 cells, where two individual foci merged to form a single spot (Fig. 2e**, Supplementary Fig. 7a and Supplementary Videos 1** and **2**). We also noted MRG splitting on some occasions (**Supplementary Fig. 7b**). Imaging TWINKLE-eGFP by SIM, we observed nucleoid “kiss-and-runs”, and nucleoid splitting, as previously described ^33^, as well as one fusion event (**Supplementary Fig. 7c, d**). In the case of fusion events, the photobleaching-corrected integrated intensity of FASTKD2-eGFP in the merged droplet was approximately the sum of the initial droplets as expected (Fig. 2f). We found this to be true for most of the fusions we measured. Notably, 75% of fusion events coincided with visible mitochondrial membrane rearrangements such as fusion, fission, or bulging (Fig. 2g).

Infoldings of the IMM called cristae densely populate the mitochondrial interior. By correlative fluorescence and electron microscopy (CLEM), we observed that displaced cristae accommodate MRGs in open spaces (Fig. 3a). Marked by fluorescence, MRGs are clearly distinguishable as round electron-dense granules, with dimensions consistent with our htSTORM data (Fig. 1 and 3a and Supplementary Fig. 8a). Noting their close proximity to the IMM, we then tested their membrane association. In mitochondria swollen by antimycin A treatment, we observed with fluorescence microscopy that most MRGs decorate the perimeter (Fig. 3b). Corroborated by similar observations in live cells (**Supplementary Fig. 8b**), this gives direct visual evidence for the previously published biochemical finding that MRG proteins co-fractionate with mitochondrial membranes, and therefore are membrane-associated ^34^.

**Fig. 3.:**
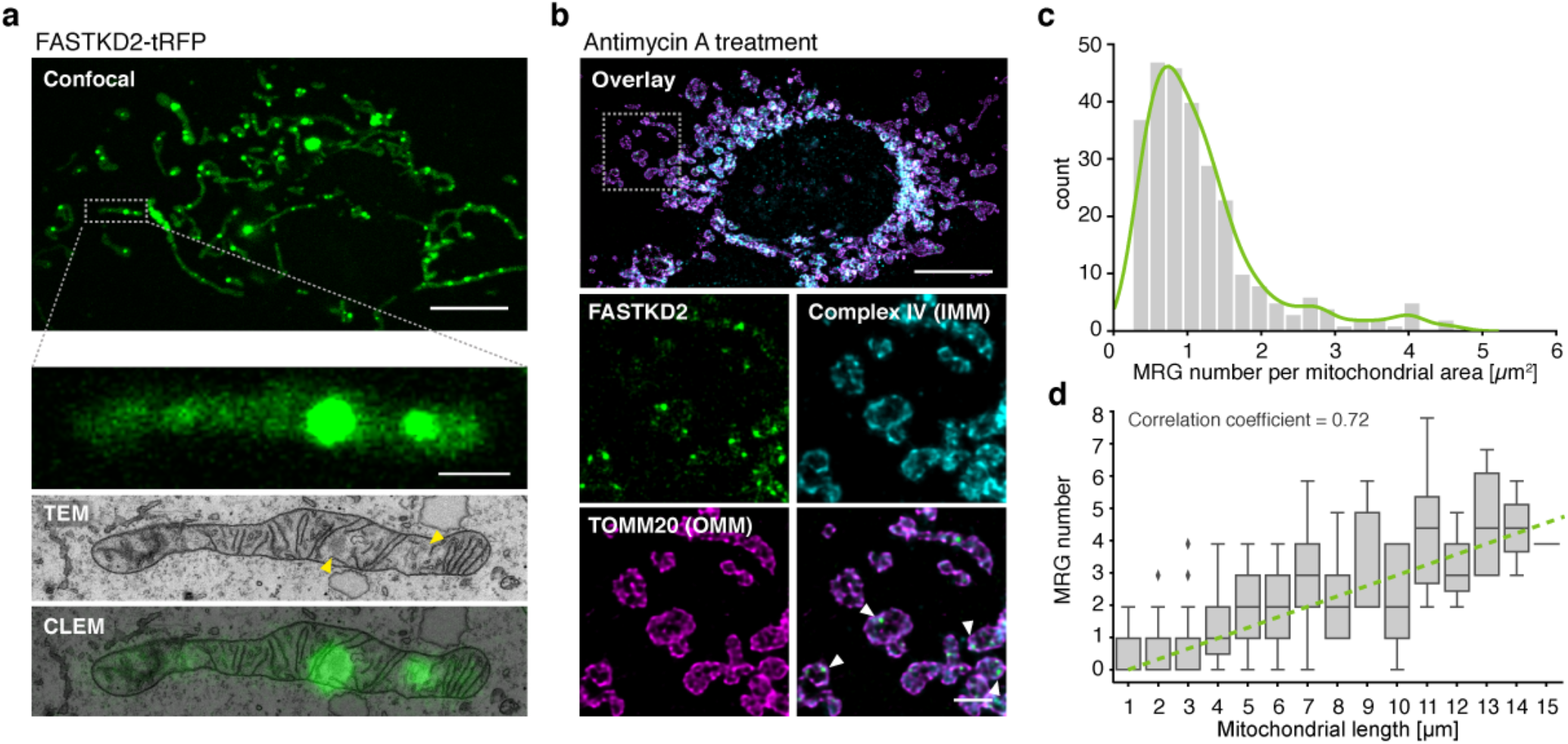
MRG positioning within mitochondria and along the mitochondrial network. **a**, Correlated transmission electron micrograph (TEM) and axially-projected confocal fluorescence image (green), with a zoom of a single mitochondrion in a FASTKD2-tRFP expressing COS-7 cell. Yellow arrowheads indicate electron densities corresponding to the MRGs. Scale bars: 10 µm (top), 1 µm (bottom). **b**, Swollen mitochondria in FASTKD2-eGFP expressing HeLa cells treated with 100 µM antimycin A for 1 hour. Inner and outer mitochondrial membranes (IMM and OMM) are immunolabeled using anti-complex IV (cyan) and anti-TOMM20 (magenta), respectively. Arrowheads in the magnified overlay highlight examples of MRGs proximal to the IMM. Scale-bars: 10 µm (top), 2 µm (zoom, bottom). **c**, Histogram of the number of MRGs per mitochondrial area. Kernel density estimate in green guides the eye. The median area occupied by a single MRG is 1 µm^2^ (± 0.8 (SD)). **d**, Dependence of MRG number on mitochondrial length. Boxplot shows a positive correlation (linear regression, green dashed line) with a correlation coefficient of 0.72. White dashed boxes indicate magnified regions.

We then assessed the distribution of MRGs along the mitochondrial network (**Supplementary Fig. 8c**). We found on average one MRG every 1 (±0.8) µm^2^ of mitochondria (n = 412), and a strong positive correlation between MRG number and mitochondrial length (correlation coefficient = 0.72, Fig. 3c, d). A non-random nucleoid distribution was found to be important for sustained inheritance of genetic material, solving the problem that given a random inheritance of nucleoids individual mitochondria would display binomial errors in partitioning ^20, 21^. Our data suggest that a similar strategy may be used for segregation of mtRNA, which could be important for maintaining steady rates of protein production in the face of mitochondrial network dynamics. To investigate the interplay between MRG positioning and mitochondrial dynamics, we inhibited mitochondrial fission by overexpression of a dominant negative mutant of the fission factor dynamin related protein 1 (Drp1), Drp1^K38A^. We observed highly elongated mitochondria with enlarged domains, as previously described and termed “mito-bulbs” ^35^. We found these domains not only contain nucleoids, but also MRGs (Fig. 4a). With super-resolved stimulated emission depletion (STED) microscopy, we discovered that mito-bulbs are better described as resembling bunches of grapes, composed of many interspersed MRGs and nucleoids, rather than as a single enlarged and coalesced structure as proposed previously ^35^ (Fig. 4b). Intrigued by the absence of MRG fusion (illustrated in Fig. 2) in such a confined space, we assessed whether MRGs may have solidified, as stress granules can ^28^. We found that fluorescence recovery of stably expressed FASTKD2-tRFP is not altered between Drp1^K38A^ overexpression and control conditions (Fig. 4c). By CLEM, we excluded the possibility that the IMM forms physical barriers between individual granules (Fig. 4d). Consistent with untreated cells (Fig. 3a), in Drp1^K38A^ expressing cells electron dense MRGs lie close to the IMM in enlarged spaces devoid of cristae. Tightly stacked cristae can be seen adjacent to mito-bulbs in affected mitochondria (Fig. 4d), rather than overlapping as previously interpreted ^35^. Knock-down of mitochondrial fusion factor Mitofusin 2 (Mfn2) evokes similar MRG clustering, further supporting a role for mitochondrial dynamics in maintaining an even distribution of MRGs along mitochondria (**Supplementary Fig. 9**).

**Fig. 4.:**
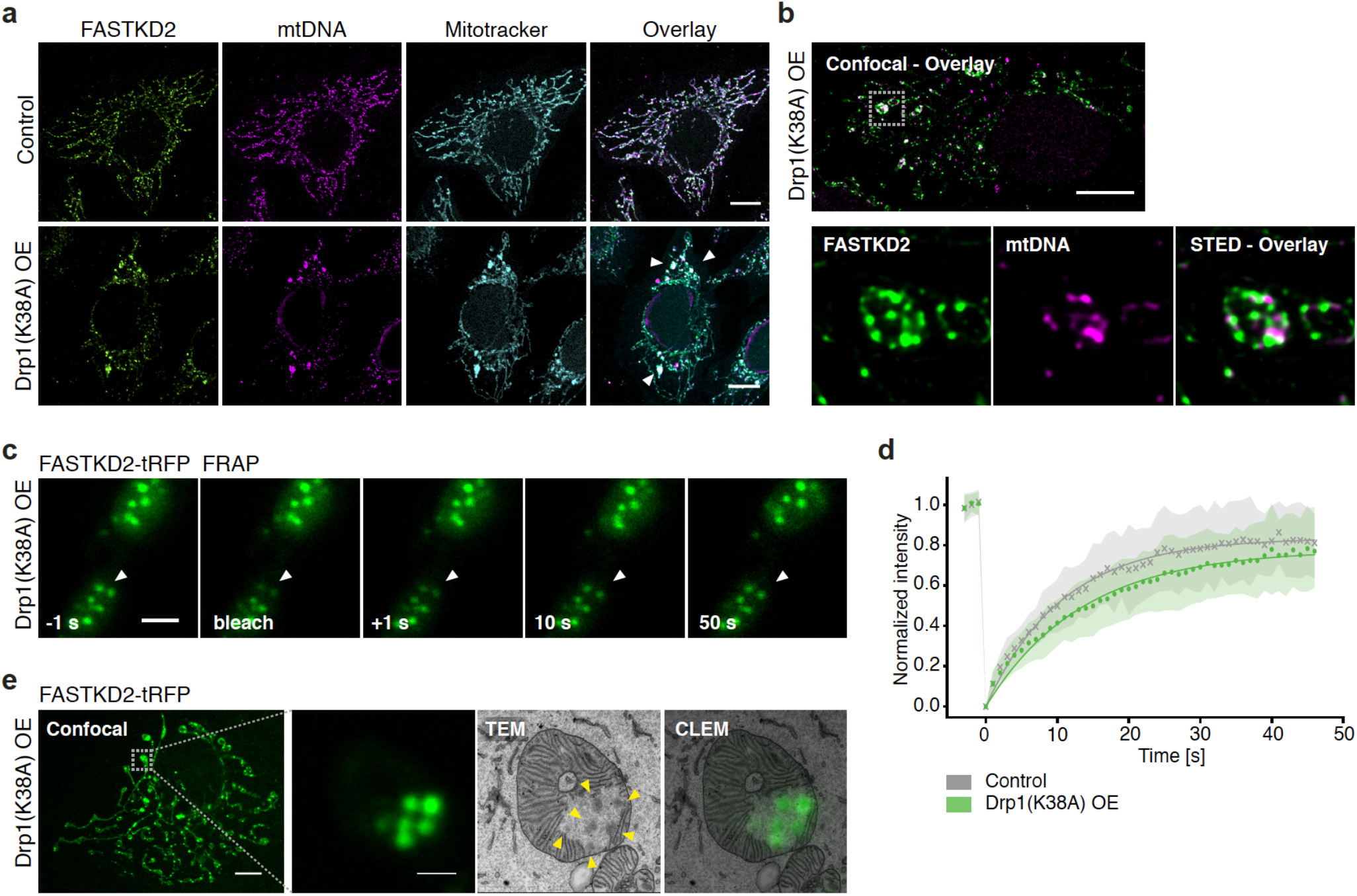
Aberrant MRG positioning in response to impaired mitochondrial fission. **a**, HeLa cells after 48 hours of Drp1^K38A^ overexpression (OE) stained using anti-FASTKD2 (MRGs, green) and anti-DNA (nucleoids, magenta) antibodies, and Mitotracker Deep Red (mitochondria, cyan). Arrowheads indicate example mito-bulbs. Scale bar: 10 µm. **b**, STED image of mito-bulbs containing MRGs (anti-FASTKD2, green) and nucleoids (anti-mtDNA, magenta) in Drp1^K38A^ overexpressing COS-7 cells. Scale bars: 10 µm (top), 2 µm (bottom, zoom). **c**, Time lapse images of mito-bulb-associated MRG FRAP experiments, in FASTKD2-tRFP (green) stably expressing COS-7 cells and transiently transfected with Drp1^K38A^ for 24 hours. Arrowheads indicate the partially bleached MRGs. Scale bar: 2 µm. **d**, Comparison of FASTKD2-tRFP FRAP between control and Drp1^K38A^ over-expressing cells. **e**, CLEM of mito-bulb-associated MRGs in COS-7 cells stably expressing FASTKD2-tRFP, fixed 24 h after Drp1^K38A^ transfection. Zoom of a single mitochondrion shows several MRGs. Overlay of MRG fluorescence and TEM (CLEM) shows electron densities corresponding to the MRGs (yellow arrowheads). Scale bars: 10µm (left), 1 µm (right). White dashed boxes indicate magnified regions.

In conclusion, our data demonstrate that MRGs are nanoscopic, internally organized liquid condensates of a characteristic size (Fig. 1 and 2). Our model proposes that condensation of mtRNA and RBPs into MRGs allows mammalian cells to regulate positioning of these components along the mitochondrial network via membrane association (Fig. 3). Mitochondrial dynamics via fission and fusion is critical for maintaining a non-random positioning of MRGs, and its perturbation disrupts their positioning while maintaining their individual stability and capacity for molecular exchange (Fig. 4). Our findings show that changes in positioning can arise, decoupled from changes in the biophysical properties of RNA and DNA sub-compartments. This insight could be important for understanding mitochondrial disorders that are reported to feature aberrant mitochondrial RNA and DNA distribution into clusters, and which imply that adequate positioning of genetic material and transcripts may be crucial for efficient ATP generation by respiration ^36^.

## Methods

### Plasmids and reagents

All cell culture reagents and chemicals were purchased from Sigma unless stated otherwise. The following plasmids were cloned in the laboratory, using pWPT lentiviral vector (Addgene #12255) as a backbone: FASTKD2-eGFP, FASTKD2-tRFP, DDX28-eGFP, ERAL1-eGFP and TWINKLE-eGFP. CFP-Drp1(K38A) plasmid was a gift from Alexander Van der Bliek. Plasmids for lentiviral production pMD2.G and psPAX2 were gifts from Didier Trono (Addgene #12259 and #12260, respectively).

### Cell culture and transfection

HeLa, COS7 and U2OS cells were cultured in Dulbecco’s modified Eagle medium (DMEM) GlutaMax supplemented with 10% heat-inactivated fetal bovine serum (FBS), 100 U/ml penicillin, 100 mg/ml streptomycin, or with 10 mM glucose and 2 mM L-glutamine in 5% CO2 at 37 °C. Cells were maintained in culture for a maximum of 20 passages and routinely assessed for mycoplasma contamination.

Plasmid transfection of cells was performed using Lipofectamine LTX (Invitrogen) or FuGENE 6 (Promega) according to manufacturer’s instructions, and cells were analyzed 24-48 hours after transfection. siRNA transfection was performed using Lipofectamine RNAi Max (Invitrogen) and cells were analyzed 72 hours after transfection.

### Stable cell lines generation

We used a second-generation lentiviral system to transform individual cell lines. In brief, gene sequences of interest were cloned into lenti-expression vector pWPT (as described above). Co-transfection of HEK293T cells was performed with packaging plasmids pMD2.G and psPAX2 using calcium phosphate precipitation. Medium containing virus was collected 48 hours after transfection and filtered using membranes with a pore size of 0.45 µm. The viral supernatant with polybrene was added to 70% confluent recipient cells, and culture medium was replaced 24 hours after infection. FACS sorting was performed to select for cells expressing GFP.

### Live-cell treatments

#### Bromouridine tagging of RNA

When bromouridine pulse assay was performed, cells were incubated with 5 mM 5-bromouridine in culture medium for 60 minutes prior to fixation, as previously described in ^3^.

#### Antimycin A treatment

For antimycin A treatment, cells were incubated with 10 mM antimycin A in Hank’s balanced salt solution (HBSS) for 60 minutes prior to fixation or live-imaging.

### Immunofluorescence and antibodies

Cells were seeded on glass coverslips and grown to a confluence of 60-80%. Following live-cell treatments if indicated, fixation of cultured cells was performed in warm 4% paraformaldehyde (PFA) in phosphate-buffer saline (PBS) for 15 min, then cells were rinsed in PBS. Cell permeabilization and blocking were done together by incubating the fixed cells in PBS containing 0.3% Triton X-100 and 1% pre-immune goat serum for 1 hour. The same buffer was used to incubate cells with the specified primary antibody (see antibody list below). After 2 hours incubation, the cells were washed in PBS and incubated with the appropriate secondary antibody conjugated with a fluorophore. Where indicated, mitochondrial network was stained before fixing the cells using Mitotracker Deep Red FM (Thermo Fisher Scientific), according to manufacturer’s instructions.

The following primary antibodies were used in this study: anti-FASTKD2 (Proteintech, 17464-1-AP), anti-bromodeoxyuridine (Roche, 11170376001), anti-GRSF1 (Sigma, HPA036985), anti-DNA (ProGen, 61014), anti-TOMM20 (Abcam, ab186734; Santa Cruz Biotech., SC-17764), anti-Complex IV (Thermo Fisher Scientific, A21348) and anti-Hsp70 (Thermo Fisher Scientific, MA3-028). Secondary antibodies were different depending on the microscopy technique applied and are detailed below.

### htSTORM

htSTORM experiments were performed as previously described ^24^, using the same hardware. Immunofluorescence was performed as described above. The following primary antibodies were combined for one-or two-color imaging, as stated in the figures: anti-FASTKD2, anti-bromodeoxyuridine, anti-GRSF1 and anti-DNA. For one color htSTORM we used Alexa Fluor 647 coupled anti-rabbit or anti-mouse secondary antibodies (Invitrogen), and respective number of foci analysed are stated in Fig. 1. To verify mitochondrial localization of analyzed foci, we co-stained the mitochondrial proteins TOMM20 or mtHSP70 using the respective primary antibodies and Alexa488 coupled secondary antibody (Invitrogen). In 2-color htSTORM, we use Alexa Fluor 647 for BrU coupled with DyLight 755 (Invitrogen) to label FASTKD2 and we analysed two cells and >30 MRGs. Before acquiring each raw STORM stack (10 ms exposure, 20,000-40,000 frames), we collected a 50 ms widefield reference image. Manually incrementing the 405-laser exposure allowed prolonged imaging. Imaging conditions (excitation illumination powers 500-1500 mW) were adjusted per sample-type. We analyzed and plotted the obtained localizations by adapting published MATLAB ^37^, and new Python scripts (see **Supplementary Fig. 1**). We calculated the full width at half maximum (FWHM) from the averaged eigenvalues as diameter for each granule. One single extreme data-point was omitted for mtDNA for creation of Fig. 1, but kept for all other analysis (incl. rendering in **Supplementary Fig. 5**). To avoid bias of non-normal distributed data, we report the median value.

### SIM live-cell microscopy

SIM was performed on a 3D NSIM Nikon microscope with a CFI Apochromat TIRF objective (100x, numerical aperture NA 1.49, Nikon). The microscope is equipped with 400 mW, 561 nm and 480 mW, 488 nm lasers (Coherent Sapphire) and a back illuminated EMCCD camera (iXon 3, Andor Technology). Live-cell imaging was performed at 37°C, using 488 and 561nm lasers for eGFP and tRFP excitation, respectively. Imaging settings were adapted to yield best image quality with minimal photo-bleaching at laser-power between 2 −10%, at 3 - 10 seconds per frame. Per field of view, 15 raw images were acquired in 3D-SIM imaging mode to ensure highest signal-to-noise ratio and resolution. Final, super-resolved SIM-images were reconstructed by the commercial Nikon NIS-Elements software and analyzed in Fiji. Opensource MicrobeJ software (www.microbej.com), originally developed for analysis of bacteria, was used for supervised automatic segmentation of mitochondria and location of their associated foci (see **Supplementary Fig. 8c**). Due to the non-Gaussian nature of the distribution (visual and Shapiro-Wilk test), we report the median value for mitochondrial occupancy. The correlation coefficient was calculated as the Pearson product moment R, with:

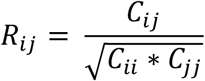

Boxplots show standard “minimum” (Q1-1.5 IQR), median and “maximum” (Q3 - 1.5 IQR) as lines, and a box from Q1 to Q3 (IQR) and “whisker-lines”, as well as outlier-points.

### FRAP and confocal microscopy

For FRAP assays, cells were seeded on coverslips and grown to 60-80% confluence. Coverslips were mounted on a Zeiss LSM 700 inverted confocal microscope with a Plan-Apochromat oil objective (63x, NA 1.40). The microscope is equipped with 488 nm and 555 nm solid-state lasers and 3 photomultipliers for simultaneous transmission and epifluorescence recording. A sliding prism and green and red bandpass filters were used to ensure clean fluorescence emission. For live assays, an Okolab stage top incubator H301 was used to maintain sample temperature at 37°C. Cells were maintained in CO_2_ independent Leibovitz L-15 medium (Gibco). For FASTKD2-(n_eGFP_ = 44, N_tRFP_ = 31, n_tRFP-Drp1KD_ = 40), ERAL1-(n = 17), and DDX28-(n = 16) FRAP the pixel size was reduced to 70 nm (Zoom 12) and line scans recorded at a pixel dwell time of 2.55 µs (maximum speed), resulting in a scan time of 97.75 ms per 128 x 128 pixel FOV. The pinhole was opened for FRAP recording. 10 x 10 pixel ROIs were manually drawn around single MRGs for FRAP and two or three pre-bleach time-points were acquired for normalization. FASTKD2-eGFP in COS7-(n = 75) & U2OS-(n = 38), and TWINKLE-eGFP (n = 56) FRAP were acquired with twice as many pixels (254×254, and 20×20 ROI) but for the same sample-size (12). A different FOV was chosen for every FRAP-experiment, multiple different cells could be imaged per sample but samples were exchanged after ∼1 h of imaging to avoid confounding effects. If MRGs had left the ROI before bleaching, the assay was aborted and a different MRG was chosen. Recovery was monitored over 50-60 s at 1 fps.

For confocal microscopy of fixed cells, samples were prepared as described above. Alexa Fluor 488, Alexa Fluor 594 or Alexa Fluor 647 secondary conjugated antibodies (Invitrogen) were used to visualize the immunolabeled targets. Imaging was performed using a Leica TCS SP8 inverted microscope with a Plan-Apochromat oil objective (63x, NA 1.4), using the Lightning mode (Leica) to generate deconvolved images. The microscope is equipped with 405 nm, 488 nm, 552 nm and 638 nm lasers.

### FRAP analysis and software

For FRAP recording of moving objects, a custom Fiji-script was co-developed with the bioimaging and optics platform at EPFL (www.biop.epfl.ch). In brief, this script “TrackFRAP” is based on the Fiji plugin TrackMate, and automatically follows the bleaching ROI during recovery. It allows the user to choose other foci as FRAP references for overall bleaching correction and outputs both a list of intensity values and metadata to allow reproducible data analysis. All tracks were manually inspected to ensure the bleached granule was recorded correctly over the full course of acquisition. If no reference granule could be tracked over the full acquisition period, the dataset was excluded from analysis. We then developed a python script to load and analyze TrackFRAP data, which we termed FRAPtrackAnalyzer [FRAPtA] and is based on the FRAPAnalyzer tool (https://omictools.com/frapanalyser-tool). Single or double exponentials were fit and plotted for each data-set, as well as used to extract recovery times.

### Correlative confocal Light and Electron Microscopy (CLEM)

Cells were seeded on a gridded coverslip (MatTek, P35-1.5-14-CGRD-D), transfected with CFP-Drp1(K38A) plasmid if applicable, and grown to 50-60% confluence. Cells were then fixed at room temperature for 1 h in fresh fixative (2% PFA, 1% glutaraldehyde in PBS 0.1M pH7.4), washed in PBS and imaged by confocal microscopy on the same day. Z-stacks were acquired of whole cells, the pinhole was closed to 0.5 AU and pixel size reduced to 50-100 nm in xy and 100-150 nm in z. Samples were then stored, overnight, in PBS at 4°C. They were then stained with osmium and potassium ferrocyanide, followed by osmium alone, each with cacodylate buffer. They were finally stained with 1% uranyl acetate, then washed in water, dehydrated through increasing concentrations of alcohol, and infiltrated with Epon resin. This was polymerized over night at 65°C. Serial, ultra-thin serial sections were then cut of the cell of interest, and the sections collected on single slot copper grids with a formvar support membrane. Images were recorded in a transmission electron microscope operating at 80kV (FEI Company, Tecnai Spirit).

### STED microscopy

For STED microscopy, samples were prepared following the immunofluorescence protocol described above. Abberior STAR 580 and Abberior STAR RED secondary antibodies (Abberior) were used in combination to label the primary antibodies. Coverslips were mounted on slides using Prolong Gold Antifade mounting agent (Thermo Fisher Scientific). Coverslips were mounted on a Leica TCS SP8 STED 3X inverted microscope equipped with an HC Plan-Apochromat glycerol motC STED W objective (93x, NA 1.30). The microscope is equipped with a white laser (470-670 nm) and 592 nm and 775 nm depletion lasers for STED. Both STED depletion lasers were set to 70% of maximum power. The pinhole was opened to 1 AU for image acquisition. Lightning mode (Leica) was used to deconvolve STED images.

## Data, code and materials availability

All imaging as well as numerical data relevant to this study are or will be publicly available on the online repository Zenodo (https://doi.org/10.5281/zenodo.3375448), or upon reasonable request. All code including adapted STORM-analysis code, TrackFRAP, FRAPtA and other python scripts for figure generation are or will be available in the online repository Github (https://github.com/TimoHenry). Both plasmids and cell lines may be available to share.

## Acknowledgments

We thank Mary-Claude Croisier of the BioEM facility at the EPFL for carrying out the electron microscopy in the CLEM, Hélène Perreten for molecular clonings, François Prodon for help with STED microscopy, and Olivier Burri and the BIOP (EPFL) for imaging support. We are grateful to Tatjana Kleele, Juliette Griffié and all members of Manley and Martinou labs for input and discussions.

## Author information

### Contributions

S.Z., T.R., J.C.M., and S.M. conceived and designed the study and wrote the manuscript. All authors reviewed and edited the manuscript. T.R., and S.Z. designed, executed, analyzed, and validated the experiments, E.C. executed and coded FRAP experiments and analysis. T.R. and S.Z. prepared the figures. S.M. and J.C.M. supervised the project.

## Ethics declarations

### Supplementary information

Supplementary Figures 1-9 and Supplementary Videos 1 and 2.

## References

1 Iborra, F. J., Kimura, H. & Cook, P. R. The functional organization of mitochondrial genomes in human cells. BMC Biol 2, 9, doi:10.1186/1741-7007-2-9 (2004).

2 Antonicka, H., Sasarman, F., Nishimura, T., Paupe, V. & Shoubridge, E. A. The mitochondrial RNA-binding protein GRSF1 localizes to RNA granules and is required for posttranscriptional mitochondrial gene expression. Cell Metab 17, 386–398, doi:10.1016/j.cmet.2013.02.006 (2013).

3 Jourdain, A. A. et al. GRSF1 regulates RNA processing in mitochondrial RNA granules. Cell Metab 17, 399–410, doi:10.1016/j.cmet.2013.02.005 (2013).

4 Handwerger, K. E., Cordero, J. A. & Gall, J. G. Cajal bodies, nucleoli, and speckles in the Xenopus oocyte nucleus have a low-density, sponge-like structure. Mol Biol Cell 16, 202–211, doi:10.1091/mbc.e04-08-0742 (2005).

5 Boeynaems, S. et al. Protein Phase Separation: A New Phase in Cell Biology. Trends Cell Biol 28, 420–435, doi:10.1016/j.tcb.2018.02.004 (2018).

6 Yamazaki, T. et al. Functional Domains of NEAT1 Architectural lncRNA Induce Paraspeckle Assembly through Phase Separation. Mol Cell 70, 1038–1053 e1037, doi:10.1016/j.molcel.2018.05.019 (2018).

7 Feric, M. et al. Coexisting Liquid Phases Underlie Nucleolar Subcompartments. Cell 165, 1686–1697, doi:10.1016/j.cell.2016.04.047 (2016).

8 Frottin, F. et al. The nucleolus functions as a phase-separated protein quality control compartment. Science, doi:10.1126/science.aaw9157 (2019).

9 Van Treeck, B. & Parker, R. Principles of Stress Granules Revealed by Imaging Approaches. Cold Spring Harb Perspect Biol 11, doi:10.1101/cshperspect.a033068 (2019).

10 Banani, S. F., Lee, H. O., Hyman, A. A. & Rosen, M. K. Biomolecular condensates: organizers of cellular biochemistry. Nat Rev Mol Cell Biol 18, 285–298, doi:10.1038/nrm.2017.7 (2017).

11 Brangwynne, C. P. et al. Germline P granules are liquid droplets that localize by controlled dissolution/condensation. Science 324, 1729–1732, doi:10.1126/science.1172046 (2009).

12 Wang, J. et al. A Molecular Grammar Governing the Driving Forces for Phase Separation of Prion-like RNA Binding Proteins. Cell 174, 688–699 e616, doi:10.1016/j.cell.2018.06.006 (2018).

13 Maharana, S. et al. RNA buffers the phase separation behavior of prion-like RNA binding proteins. Science 360, 918–921, doi:10.1126/science.aar7366 (2018).

14 Langdon, E. M. et al. mRNA structure determines specificity of a polyQ-driven phase separation. Science 360, 922–927, doi:10.1126/science.aar7432 (2018).

15 Van Treeck, B. & Parker, R. Emerging Roles for Intermolecular RNA-RNA Interactions in RNP Assemblies. Cell 174, 791–802, doi:10.1016/j.cell.2018.07.023 (2018).

16 Alberti, S., Gladfelter, A. & Mittag, T. Considerations and Challenges in Studying Liquid-Liquid Phase Separation and Biomolecular Condensates. Cell 176, 419–434, doi:10.1016/j.cell.2018.12.035 (2019).

17 Jourdain, A. A., Boehm, E., Maundrell, K. & Martinou, J. C. Mitochondrial RNA granules: Compartmentalizing mitochondrial gene expression. J Cell Biol 212, 611–614, doi:10.1083/jcb.201507125 (2016).

18 Pearce, S. F. et al. Regulation of Mammalian Mitochondrial Gene Expression: Recent Advances. Trends Biochem Sci 42, 625–639, doi:10.1016/j.tibs.2017.02.003 (2017).

19 Pernas, L. & Scorrano, L. Mito-Morphosis: Mitochondrial Fusion, Fission, and Cristae Remodeling as Key Mediators of Cellular Function. Annu Rev Physiol 78, 505–531, doi:10.1146/annurev-physiol-021115-105011 (2016).

20 Jajoo, R. et al. Accurate concentration control of mitochondria and nucleoids. Science 351, 169–172, doi:10.1126/science.aaa8714 (2016).

21 Lewis, S. C., Uchiyama, L. F. & Nunnari, J. ER-mitochondria contacts couple mtDNA synthesis with mitochondrial division in human cells. Science 353, aaf5549, doi:10.1126/science.aaf5549 (2016).

22 Ghezzi, D. et al. FASTKD2 nonsense mutation in an infantile mitochondrial encephalomyopathy associated with cytochrome c oxidase deficiency. Am J Hum Genet 83, 415–423, doi:10.1016/j.ajhg.2008.08.009 (2008).

23 Yoo, D. H. et al. Identification of FASTKD2 compound heterozygous mutations as the underlying cause of autosomal recessive MELAS-like syndrome. Mitochondrion 35, 54–58, doi:10.1016/j.mito.2017.05.005 (2017).

24 Douglass, K. M., Sieben, C., Archetti, A., Lambert, A. & Manley, S. Super-resolution imaging of multiple cells by optimised flat-field epi-illumination. Nat Photonics 10, 705–708, doi:10.1038/nphoton.2016.200 (2016).

25 Brown, T. A. et al. Superresolution fluorescence imaging of mitochondrial nucleoids reveals their spatial range, limits, and membrane interaction. Mol Cell Biol 31, 4994–5010, doi:10.1128/MCB.05694-11 (2011).

26 Kukat, C. et al. Super-resolution microscopy reveals that mammalian mitochondrial nucleoids have a uniform size and frequently contain a single copy of mtDNA. Proc Natl Acad Sci U S A 108, 13534–13539, doi:10.1073/pnas.1109263108 (2011).

27 Alan, L., Spacek, T. & Jezek, P. Delaunay algorithm and principal component analysis for 3D visualization of mitochondrial DNA nucleoids by Biplane FPALM/dSTORM. Eur Biophys J 45, 443–461, doi:10.1007/s00249-016-1114-5 (2016).

28 Jain, S. et al. ATPase-Modulated Stress Granules Contain a Diverse Proteome and Substructure. Cell 164, 487–498, doi:10.1016/j.cell.2015.12.038 (2016).

29 Zaganelli, S. et al. The Pseudouridine Synthase RPUSD4 Is an Essential Component of Mitochondrial RNA Granules. J Biol Chem 292, 4519–4532, doi:10.1074/jbc.M116.771105 (2017).

30 Tu, Y. T. & Barrientos, A. The Human Mitochondrial DEAD-Box Protein DDX28 Resides in RNA Granules and Functions in Mitoribosome Assembly. Cell Rep 10, 854–864, doi:10.1016/j.celrep.2015.01.033 (2015).

31 Farge, G. et al. The N-terminal domain of TWINKLE contributes to single-stranded DNA binding and DNA helicase activities. Nucleic Acids Res 36, 393–403, doi:10.1093/nar/gkm1025 (2008).

32 Wheeler, J. R., Matheny, T., Jain, S., Abrisch, R. & Parker, R. Distinct stages in stress granule assembly and disassembly. Elife 5, doi:10.7554/eLife.18413 (2016).

33 Garrido, N. et al. Composition and dynamics of human mitochondrial nucleoids. Mol Biol Cell 14, 1583–1596, doi:10.1091/mbc.e02-07-0399 (2003).

34 Jourdain, A. A. et al. A mitochondria-specific isoform of FASTK is present in mitochondrial RNA granules and regulates gene expression and function. Cell Rep 10, 1110–1121, doi:10.1016/j.celrep.2015.01.063 (2015).

35 Ban-Ishihara, R., Ishihara, T., Sasaki, N., Mihara, K. & Ishihara, N. Dynamics of nucleoid structure regulated by mitochondrial fission contributes to cristae reformation and release of cytochrome c. Proc Natl Acad Sci U S A 110, 11863–11868, doi:10.1073/pnas.1301951110 (2013).

36 Durigon, R. et al. LETM1 couples mitochondrial DNA metabolism and nutrient preference. EMBO Mol Med 10, doi:10.15252/emmm.201708550 (2018).

37 Sieben, C., Banterle, N., Douglass, K. M., Gonczy, P. & Manley, S. Multicolor single-particle reconstruction of protein complexes. Nat Methods 15, 777–780, doi:10.1038/s41592-018-0140-x (2018).

