## Supplementary material for "Mitochondrial RNA granules are fluid condensates, positioned by membrane dynamics": SI

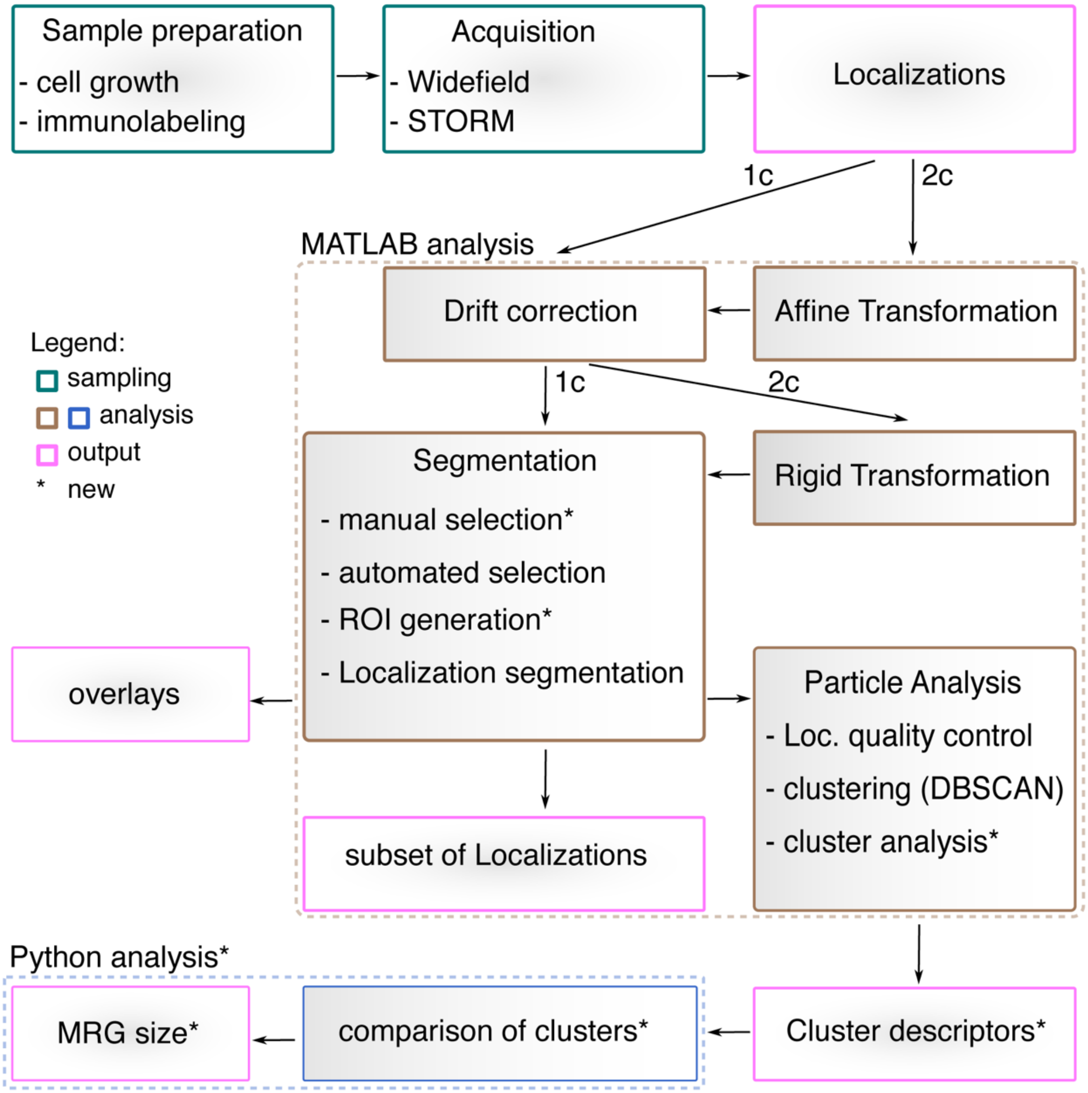


**Supplementary Fig. 1: Workflow for granule analysis using htSTORM.**
htSTORM acquisition and analysis were performed by adapting previously published workflows and analysis scripts [24,37]. New, previously unpublished parts of the analysis are highlighted by asterisks.

Supplementary figure 2 - Rey, Zaganelli et al.

FASTKD2 clusters after DBSCAN clustering analysis (n=326)

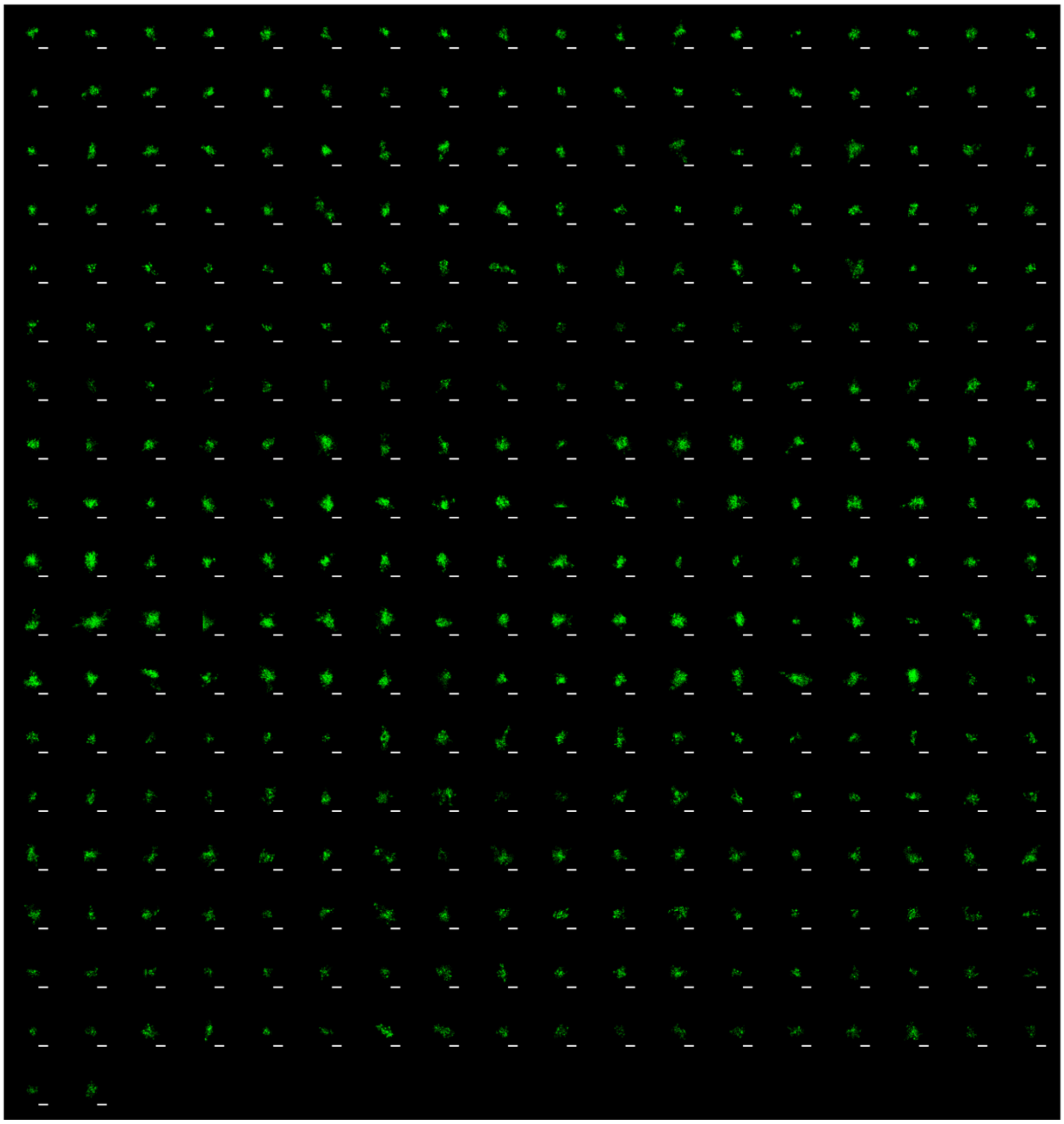

**Supplementary Fig. 2: Montage of FASTKD2 granules imaged with htSTORM (one color).** Montage of 326 granules after DBSCAN clustering of MRGs immunolabeled using anti-FASTKD2 antibody. These data correspond to those presented in **Fig. 1b**. Scale bars: 200 nm.

Supplementary figure 3 - Rey, Zaganelli et al.

GRSF1 clusters after DBSCAN clustering analysis (n=361)

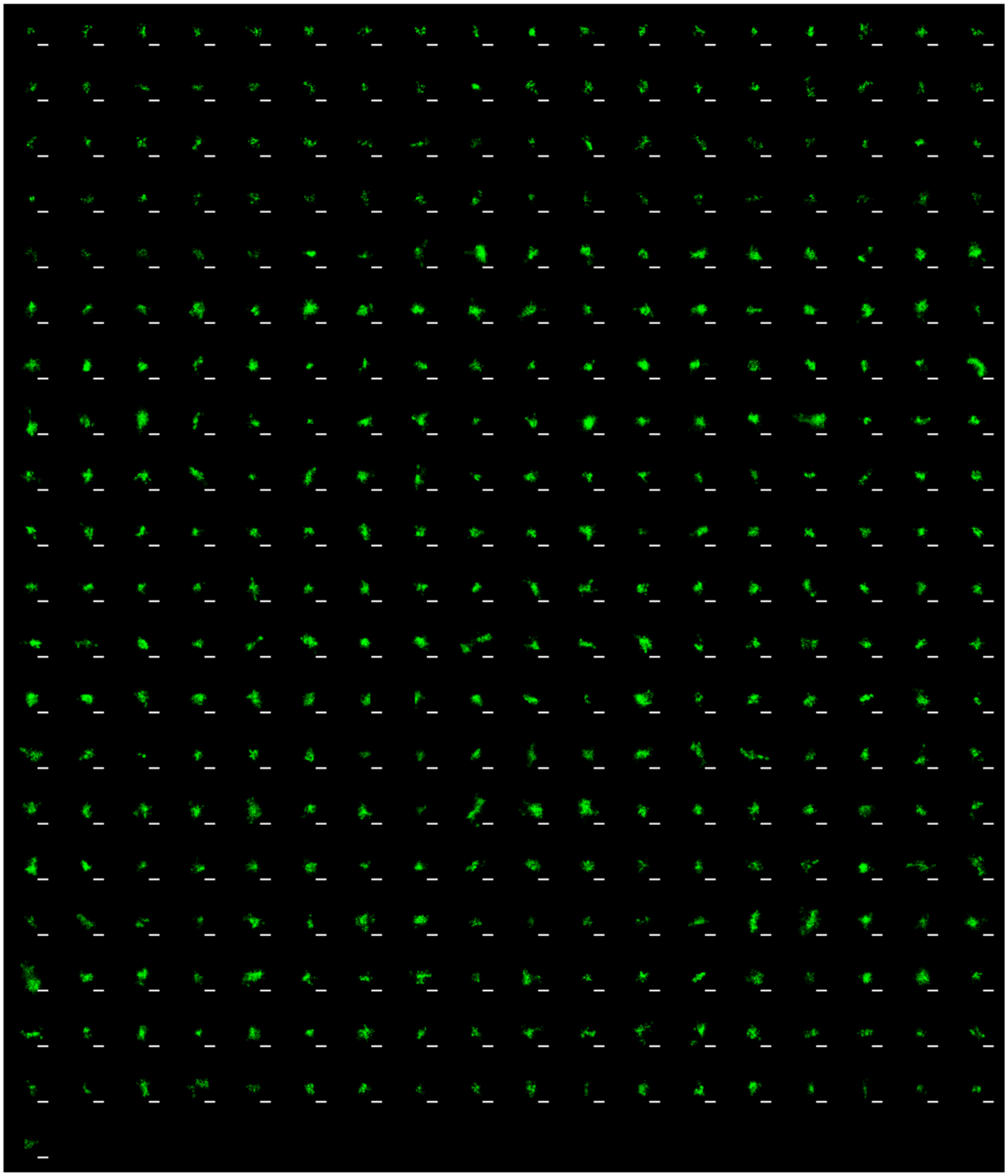

**Supplementary Fig. 3: Montage of GRSF1 granules imaged with htSTORM (one color).** Montage of 361 granules after DBSCAN clustering of MRGs immunolabeled using anti-GRSF1 antibody. These data correspond to those presented in **Fig. 1b**. Scale bars: 200 nm.

Supplementary figure 4 - Rey, Zaganelli et al.

mtRNA (BrU) clusters after DBSCAN clustering analysis (n=254)

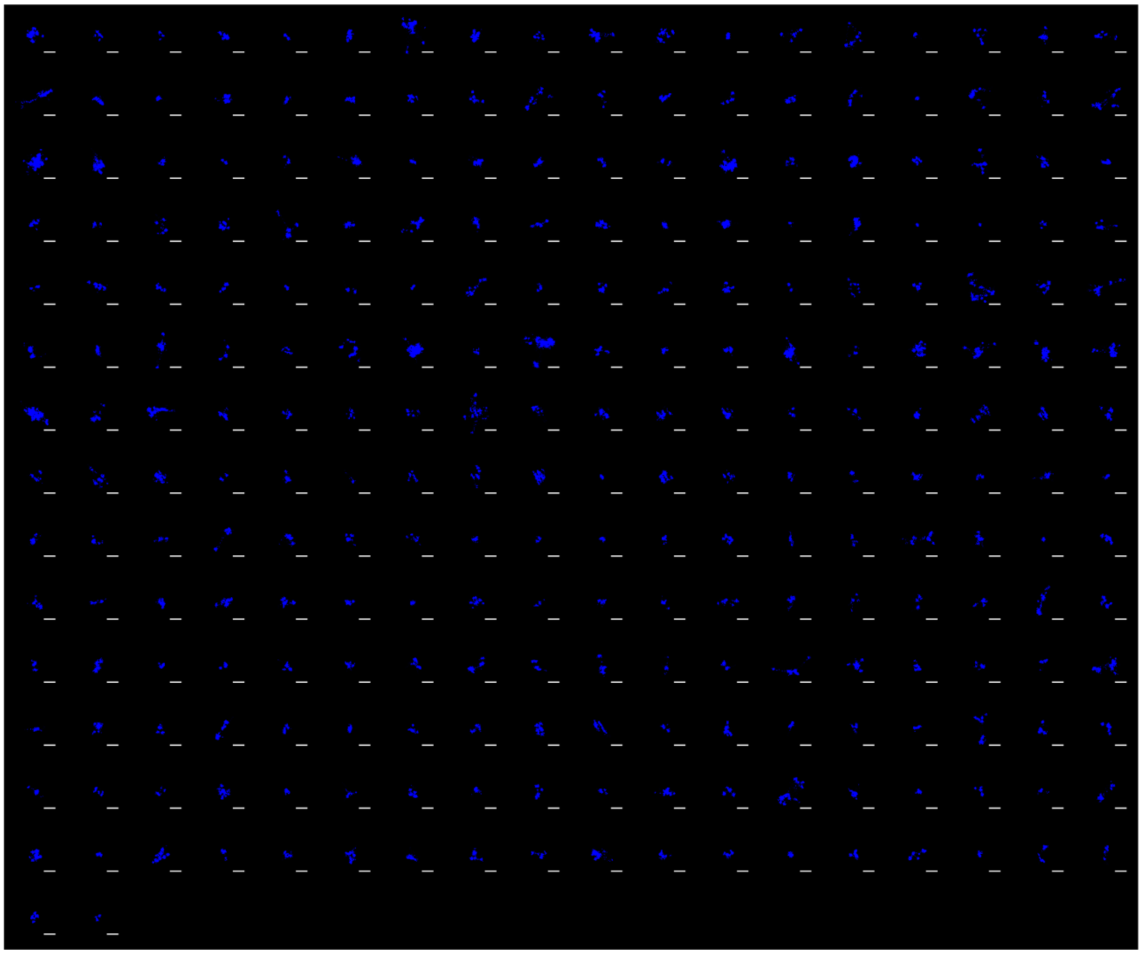

**Supplementary Fig. 4: Montage of mtRNA granules imaged with htSTORM (one color).** Montage of 254 granules after DBSCAN clustering of MRGs immunolabeled using anti-BrdU antibody, following a 1h bromouridine incubation of live cells prior to fixation. These data correspond to those presented in **Fig. 1b**. Scale bars: 200 nm.

Supplementary figure 5 - Rey, Zaganelli et al.

mtDNA clusters after DBSCAN clustering analysis (n=431)

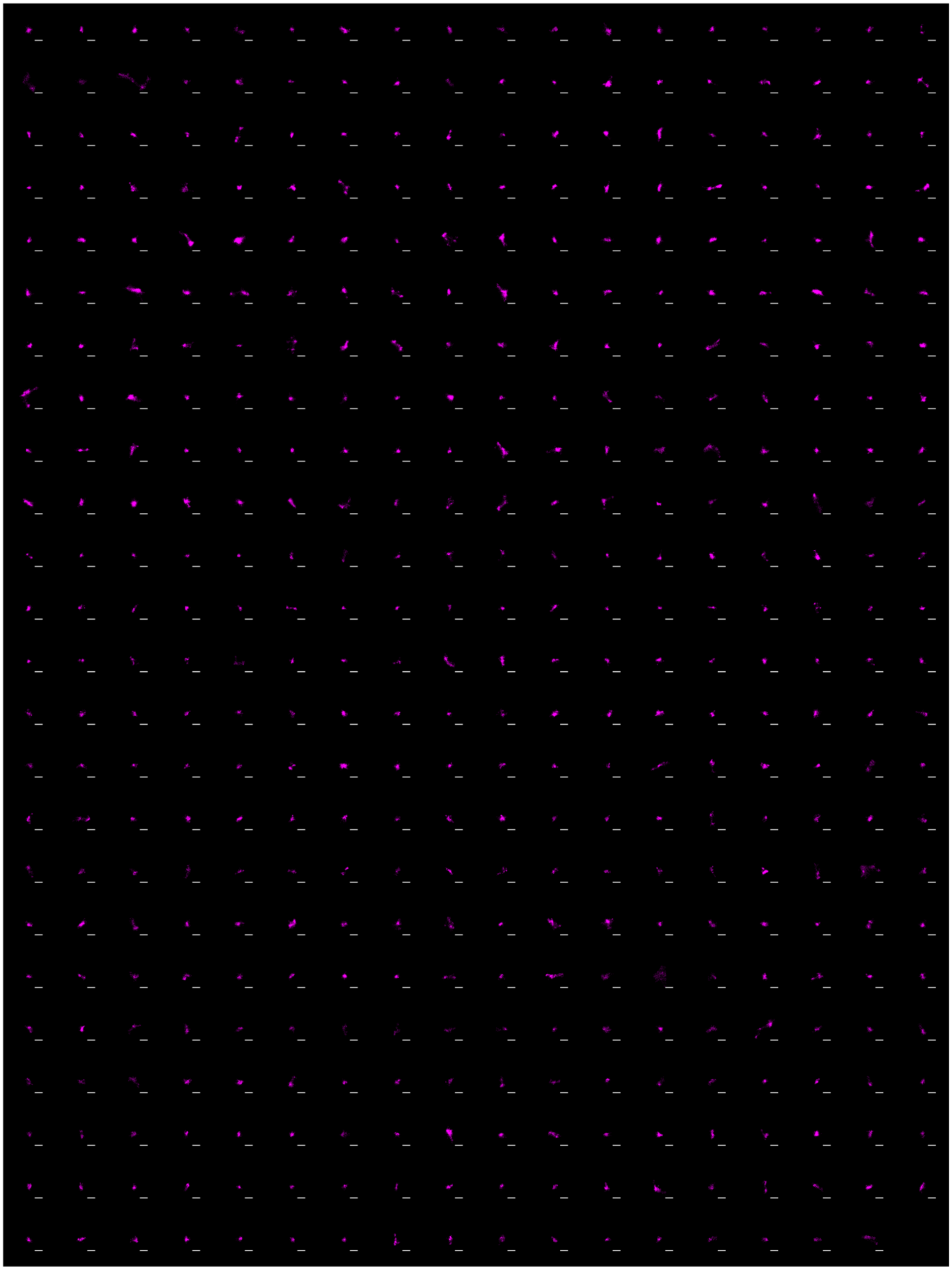

**Supplementary Fig. 5: Montage of mtDNA granules imaged with htSTORM (one color).** Montage of 431 granules after DBSCAN clustering of nucleoids immunolabeled using anti-DNA antibody. These data correspond to those presented in **Fig. 1b**. Scale bars: 200 nm.

Supplementary figure 6 - Rey, Zaganelli et al.

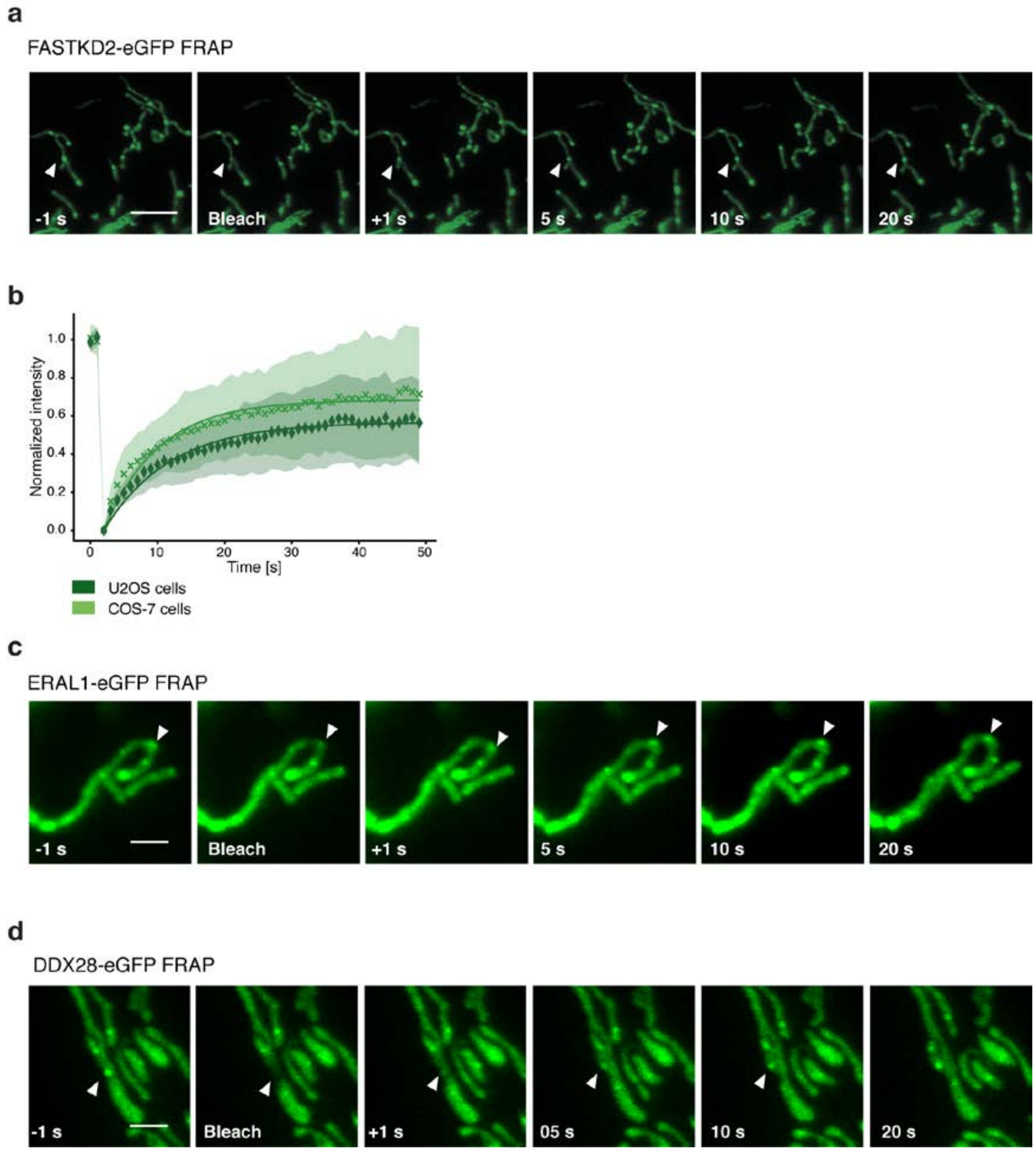


**Supplementary Figure 6 FRAP of MRG-associated proteins.**

**a,** Representative time lapse images of MRGs FRAP experiments in U2OS cells stably expressing FASTKD2-eGFP (green). White arrowheads indicate the photobleached structures. Scale bar: 5 µm. **b,** FRAP analysis of FASTKD2-eGFP in U2OS and COS-7 cells. Symbols in the graph represent mean data points. Single exponential fits (lines) and standard deviations for each time point (shaded area) are shown. **c, d,** Representative time lapse images of ERAL1- and DDX28-eGFP FRAP experiments in COS-7 cells. White arrowheads indicate the photobleached structures. These images correspond to the data plotted in **Fig. 2c** Scale bar: 2 µm.

Supplementary figure 7 - Rey, Zaganelli et al.

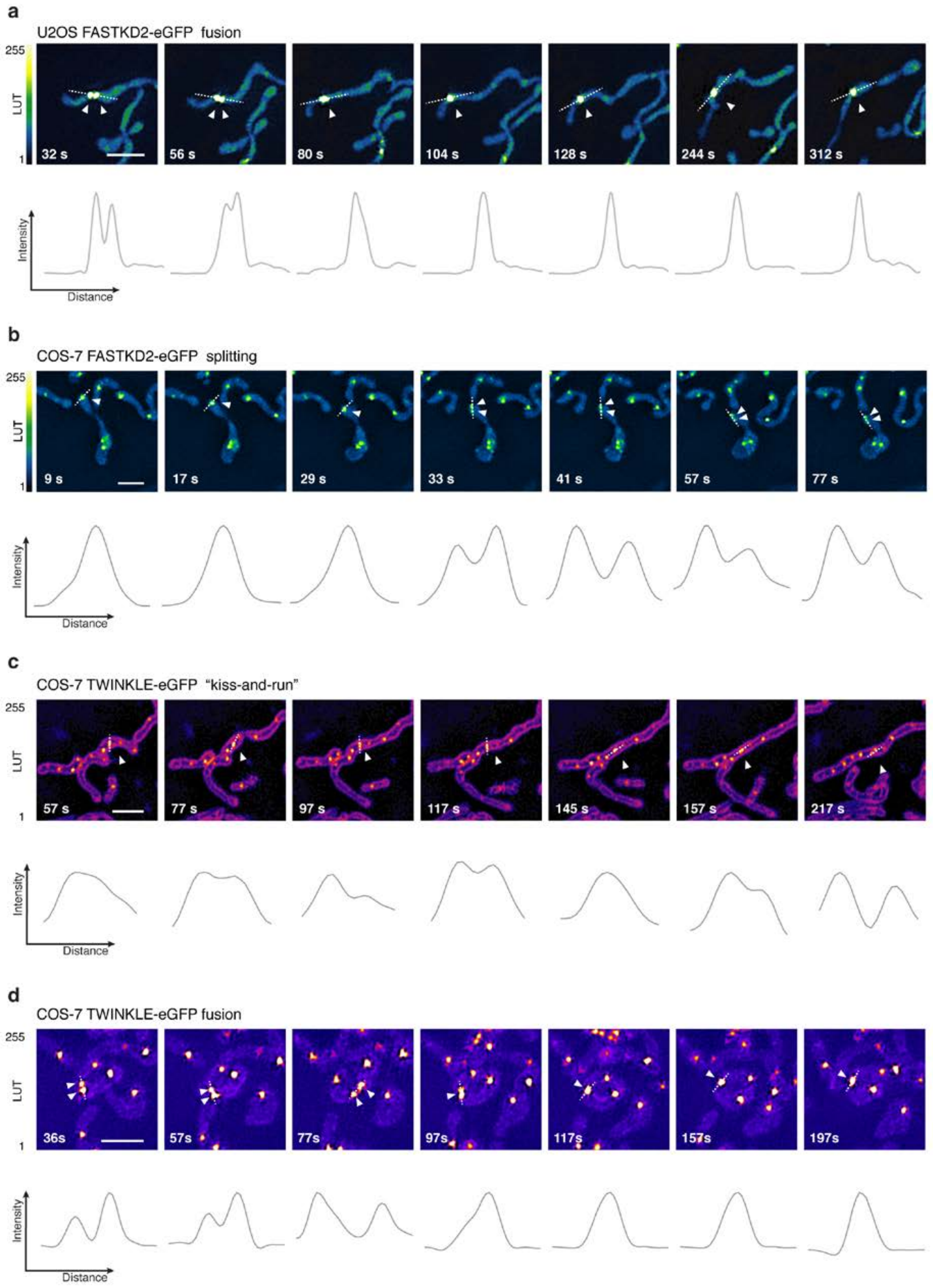

**Supplementary Fig. 7: MRG and nucleoid dynamics in live cells.**

**a** Representative time lapse images of an MRG fusion event in a live U2OS cell, monitored by SIM. MRGs are visualized by stable expression of FASTKD2-eGFP. **b,** Representative time lapse images of an MRG splitting event in a live COS-7 cell, monitored by SIM. MRGs are visualized by stable expression of FASTKD2-eGFP. **c, d,** Representative time lapse images of nucleoid "kiss-and-run" and splitting events, respectively, in COS-7 cells, monitored by SIM. Nucleoids are visualized by transient expression of TWINKLE-eGFP and mitochondrial outlines are highlighted by TOMM20-eGFP expression. **a-d** Cells were imaged at 1/5 Hz. White arrowheads indicate the dynamic events. Dashed lines indicate the segments used to measure the intensity (grey values) represented in the plots below. Scale bars: 2 µm.

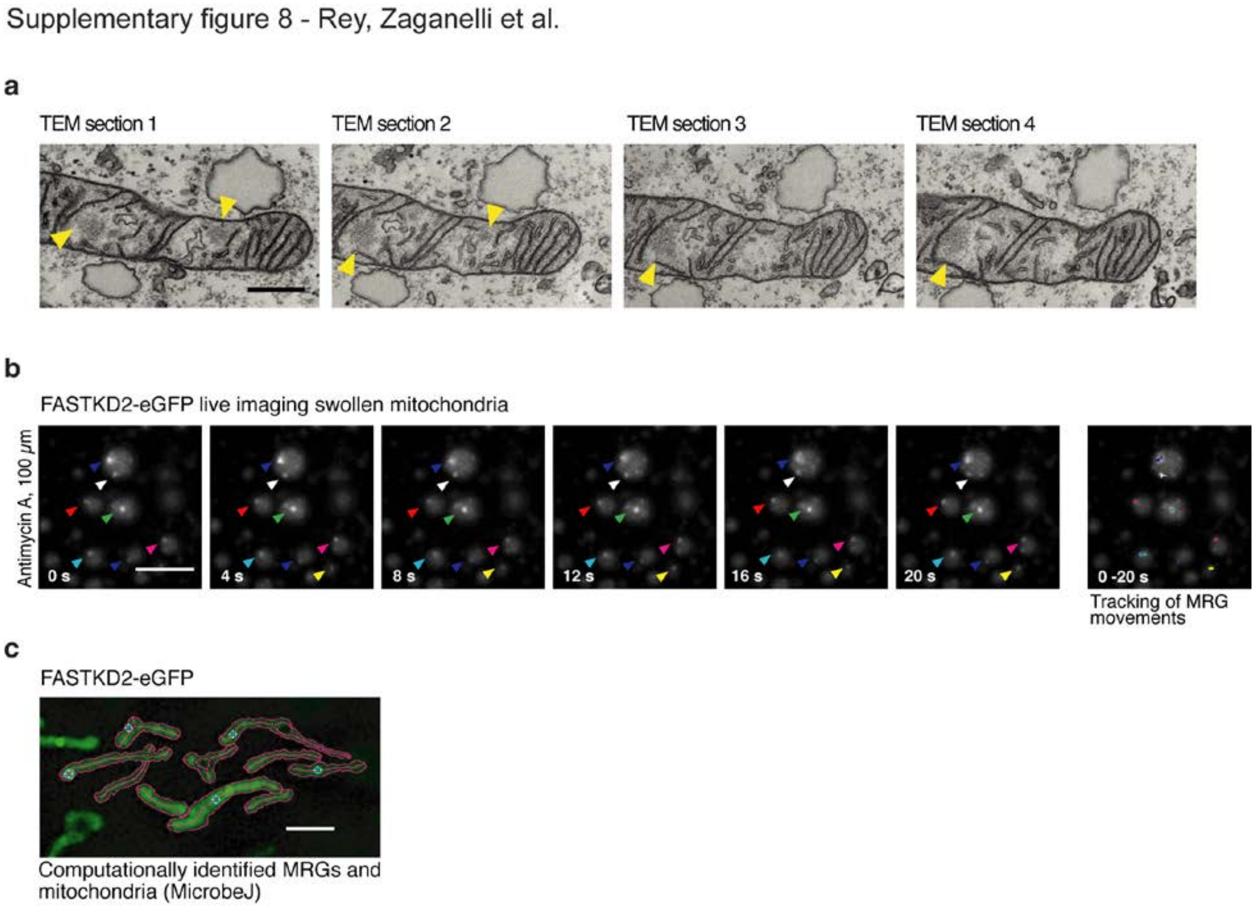


**Supplementary Fig. 8: MRGs are associated with the IMM and evenly distributed along the mitochondrial network.**

**a** TEM highlights corresponding to the data shown in **Fig. 3a**. Different contiguous TEM microtome sections show electron densities corresponding to the MRGs visualized by fluorescence microscopy (yellow arrowheads). Scale bar: 500 nm. **b** Representative time lapse images of swollen mitochondria. HeLa cells were treated with 100 µM antimycin A to induce mitochondrial swelling during 1 hour prior to live-cell imaging. Movements of single MRGs over time were tracked using the Manual Tracking ImageJ plugin (right image). Arrowheads indicate the MRGs analyzed. Scale-bar: 2 µm. **c** Semi-automated mitochondria segmentation and MRG-association with their parent organelle with the ImageJ plugin, MircrobeJ.

Supplementary figure 9 - Rey, Zaganelli et al.

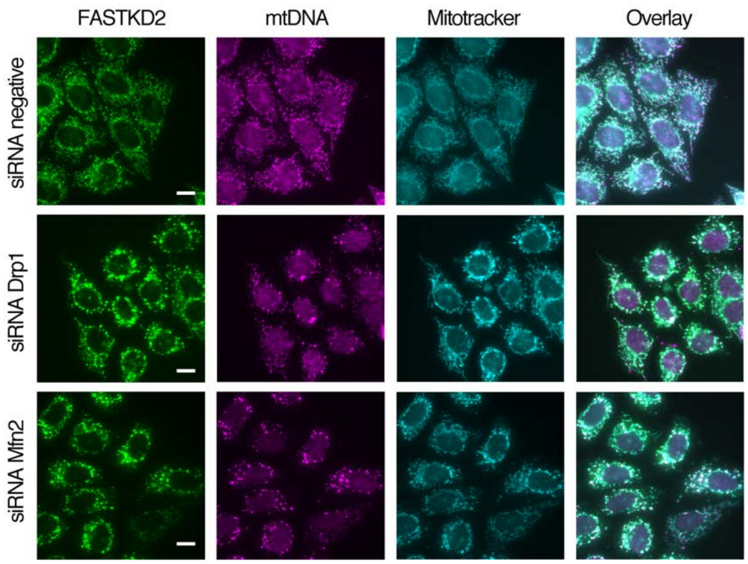


**Supplementary Fig. 9: Silencing of Drp1 and Mfn2 have similar effects on MRG and nucleoid positioning.**

Representative widefield images of HeLa cells silenced using siRNAs against Drp1 and Mfn2. Cells were fixed after 72 hours of silencing. A negative siRNA was used as a control. MRGs and nucleoids were immunolabeled using anti-FASTKD2 (green) and anti-DNA (magenta), respectively. Mitochondria were labeled using Mitotracker Deep Red staining (cyan). Scale-bar: 10 µm.

**Supplementary Video 1. Fusion of MRGs in live COS-7 cells.**

SIM of an MRG fusion event in cells stably expressing FASTKD2-eGFP (green). Movie corresponding to the data shown in **Fig. 2e**. Scale bar: 2 µm.

**Supplementary Video 2. Fusion of MRGs in live U2OS cells.**

SIM of an MRG fusion event in cells stably expressing FASTKD2-eGFP (green). Movie corresponding to the data shown in **Supplementary Fig. 7a**. Scale bar: 2 µm.
